# Full three-dimensional imaging deep through multicellular thick samples with subcellular resolution by structured illumination microscopy and adaptive optics

**DOI:** 10.1101/2020.04.15.043026

**Authors:** Ruizhe Lin, Edward T. Kipreos, Jie Zhu, Chang Hyun Khang, Peter Kner

## Abstract

Structured Illumination Microscopy enables live imaging with resolutions of ~120 nm. Unfortunately, optical aberrations can lead to loss of resolution and artifacts in Structured Illumination Microscopy rendering the technique unusable in samples thicker than a single cell. Here we report on the combination of Adaptive Optics and Structured Illumination Microscopy enabling imaging with 140 nm lateral and 585 nm axial resolution in tissue culture cells, *C. elegans*, and rice blast fungus. We demonstrate that AO improves resolution and reduces artifacts, making full 3D SIM possible in thicker samples.

## Introduction

Fluorescence microscopy is a critical tool for biological discovery, and recent advances in fluorescence microscopy have led to significant advances in biology [1-3]. Different techniques have been developed to image below the diffraction limit including structured illumination microscopy (SIM) [4, 5], stimulated emission depletion microscopy (STED) [6, 7], photoactivated localization microscopy (PALM) [8, 9], and stochastic optical reconstruction microscopy (STORM) [10-12]. Among these super-resolution (SR) techniques, SIM stands out for its compatibility with live-cell imaging [3, 13]. As a widefield-based method, with low phototoxicity, SIM achieves lateral resolution of ~120 nm and axial resolution of ~ 300 nm [14, 15], compared to that of ~250 nm in conventional widefield microscopes. And SIM requires significantly fewer image acquisitions than required for STORM or PALM for every SR image reconstruction allowing for acquisition rates fast enough for live cell imaging [5, 13]. Moreover, three dimensional SIM (3D-SIM) provides optical sectioning [16], which eliminates out-of-focus light [17, 18]. While STED can also be used for live imaging, STED typically requires higher laser intensities and cannot image large fields of view as rapidly.

Unfortunately, for all fluorescence microscopes, the existence of aberrations results in image deterioration [19, 20]. Aberrations are especially significant when imaging deep into live organisms, and SR methods are generally more sensitive to aberrations [21]. In SIM, aberrations in the widefield optical transfer function (OTF) not only lead to noise, artifacts and loss of resolution in the SIM image [22] but can also prevent the SIM reconstruction algorithm from finding a solution at all. The aberrations can be classified into two categories, namely system aberrations and sample induced aberrations. System aberrations are due to defects of optical elements or the imperfect optical alignment in the microscope system and directly affect the point spread function (PSF) of the microscope. Sample-induced aberrations are caused by the optical properties of the biological sample including the difference in refractive index between the sample and the surrounding media, and the refractive index variations within the sample itself [23, 24]. Due to the diverse nature of biological samples, sample-induced aberrations are case-specific, and frequently the aberrations are spatially variant within the sample.

To correct the aberrations and restore the image quality, adaptive optics (AO) provides a feasible solution [25-27] and has been applied in several microscopy systems [28-31]. Direct wavefront sensing employs a wavefront sensor to provide a direct instantaneous measurement of the wavefront [32, 33]. Its fast response allows the deformable mirror (DM) to be adjusted for every image frame when imaging dynamic processes in live samples. Turcotte et al. [34] reported a combination of 2D SR-SIM (increased lateral resolution and optical sectioning) and direct wavefront sensing AO to image the dynamics of dendrites and dendritic spines in the living mouse brain *in vivo.* However, the direct wavefront sensing method requires a dedicated wavefront sensing system (e.g. a Shack–Hartmann wavefront sensor), and an isolated guide-star (e.g. two photon-induced fluorescent guide star [35, 36] or fluorescent protein guide-star [37, 38]) which increases the system complexity and cost. Moreover, when the wavefront sensing system fails to form an identifiable image of the guide-star due to large aberrations or highly scattering medium, it loses its efficiency dramatically. Alternatively, a sensorless model-based AO method is more resistant to scattering and can be operated upon highly aberrated images [39-41]. It relies on a series of artificial aberration trials and an image quality metric function. The optimization is conducted iteratively by maximizing the metric value. This AO method has been applied to optical sectioning SIM [16, 42] and two-dimensional SR-SIM [22]. Žurauskas et al. [43] reported on an AO structured illumination system with a sensorless AO method that ensures adequate sampling of high frequency information. The samples being imaged are flat cells cultured and mounted on glass, which don’t present the large aberrations existing in thick multicellular organisms, and the final image reconstruction is not fully extended to three-dimensional SIM.

In this article, we demonstrate the application of AO to 3D-SIM, achieving a resolution of 143 nm laterally and 585 nm axially (emission wavelength at 670 nm, NA of 1.2), along with optical sectioning capability, compared to a resolution of 279 nm laterally and 930 nm axially in widefield imaging. The sensorless, image-based AO method is shown to be capable of correcting system aberrations and sample induced aberrations in a variety of samples with minimal photobleaching. We show that using structured illumination and a spatial frequency based metric function helps the AO correction performance in thick live organisms. To demonstrate AO-3DSIM, we took full three-dimensional images of α-TN4 cells, fluorescent beads under the body of a *C. elegans* worm, GFP labeled axons in *C. elegans*, and the GFP labelled endoplasmic reticulum of rice blast fungus inside rice plant cells. The final 3D images clearly reveal the object of interest in all three dimensions, and the application of AO yields a significant improvement in the image. Image signal to noise ratio (SNR) is significantly increased with the application of AO, and fine structures which are not identifiable when aberrations are present, become well resolved and identifiable.

## Methods

### Three-dimensional Structured Illumination

We follow the method for three-dimensional Structured Illumination Microscopy developed by Gustafsson et al [18]. To resolve the pattern from the raw data and reconstruct the super resolution image, fifteen raw images are acquired for each plane, which consist of three pattern orientations (0°, 120°, 240°), and five equally distributed phases 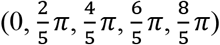 for each angle. The three-dimensional data is acquired as a sequence of two-dimensional images as the sample is stepped through the focal plane. The final three-dimensional image is obtained through a linear recombination of all post-processed and frequency shifted images 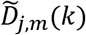, using a generalized Wiener filter:

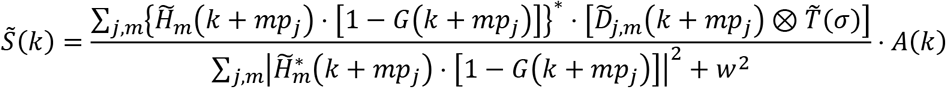

where *w*^2^ is the Wiener parameter, *mp_j_* denotes *m*^th^ order frequency component at the *j*th pattern orientation. is the Fourier Transform of the data for the m^th^ order and j^th^ angle. 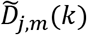 is the Optical Transfer Function for the *m*^th^ order; the OTF for the first order consists of two copies of the zeroth order OTF shifted in the axial direction. *G*(*k* + *mp_j_*) is a top-hat notch filter with a diameter of 9 pixels which suppresses the overemphasized zero frequency of each OTF copy, removing the residual stripe artifacts and enhancing the optical sectioning [44, 45]. *T*(*γ*) is a Tukey window function (*γ* = 0.08) that is used to eliminate the edge effect in the Fast Fourier transform (FFT) due to edge discontinuities. *A*(*k*) is the apodization function to cancel the ringing of high frequency spectral components caused by the Wiener filter. We used a modified three dimensional triangle function *A*(*k*) = [(1 − a_x_y · k_x_y/k_cutoff_xy_) · (1 − a_z_ · k_z_/k_cutoff_z_)]^n^, where the cutoff frequencies *k_cutoff_xy_* = 4 · *NA*/*λ, k_cutoff_z_* = *NA*^2/^*λ*. The parameters *a_xy_* ∈ [0.5,1], *a_z_* ∈ [0.5,1] and *n* ∈ [0.5,2] are tuned empirically to balance the resolution loss and the artifact elimination. All functions are defined in a three dimensional volume with the size of 1024 × 1024 × *n_z_*, in which *n_z_* is determined by the number of slices in the z-stack.

### AO

The trial aberration modes are represented by the Zernike modes, which are a complete, orthogonal set of eigenfunctions defined over the unit circle [46]. As is well known, the low-order Zernike modes correspond to the typical aberrations of an optical system [47]. In the limit of small aberrations, each of the orthogonal Zernike modes affects the image independently from others, so the final correction is built on a linear summation of all modes to be corrected. In most cases, we correct Zernike modes 4, 5, 6, 7 and 10, which are responsible for astigmatism, coma, and spherical aberration.

The simulation results in Fig. 1 shows the impact of three basic aberration modes (Astigmatism, Coma and Spherical aberration – Fig. 1b) on the 3D-SIM spatial images, Fig. 1d, and effective OTFs, Fig. 1e. As they do in the widefield image, the aberrations result in additional intensity structure around the PSF which will affect the signal to noise. The aberrations also strongly weaken the effective OTF at higher frequencies due to both the aberrations reducing the widefield OTF strength and the reduction in strength of the higher orders. From the graphs in Fig. 1f, we can see the decrease of computed pattern amplitudes as the aberration strength increases. The amplitude of some patterns decreases by more than 300% when the pattern orientation partially coincides with the orientation of the aberration modes. It is worth mentioning that we run the simulations with preset pattern frequency and phase, so we achieve good parameter fitting when running the reconstruction algorithm on aberrated images. However, in real cases, the drop of pattern amplitudes could result in inaccurate computation of the pattern frequency and phase, which would introduce more serious artifacts than what we see in Fig. 1.

**Fig 1.**
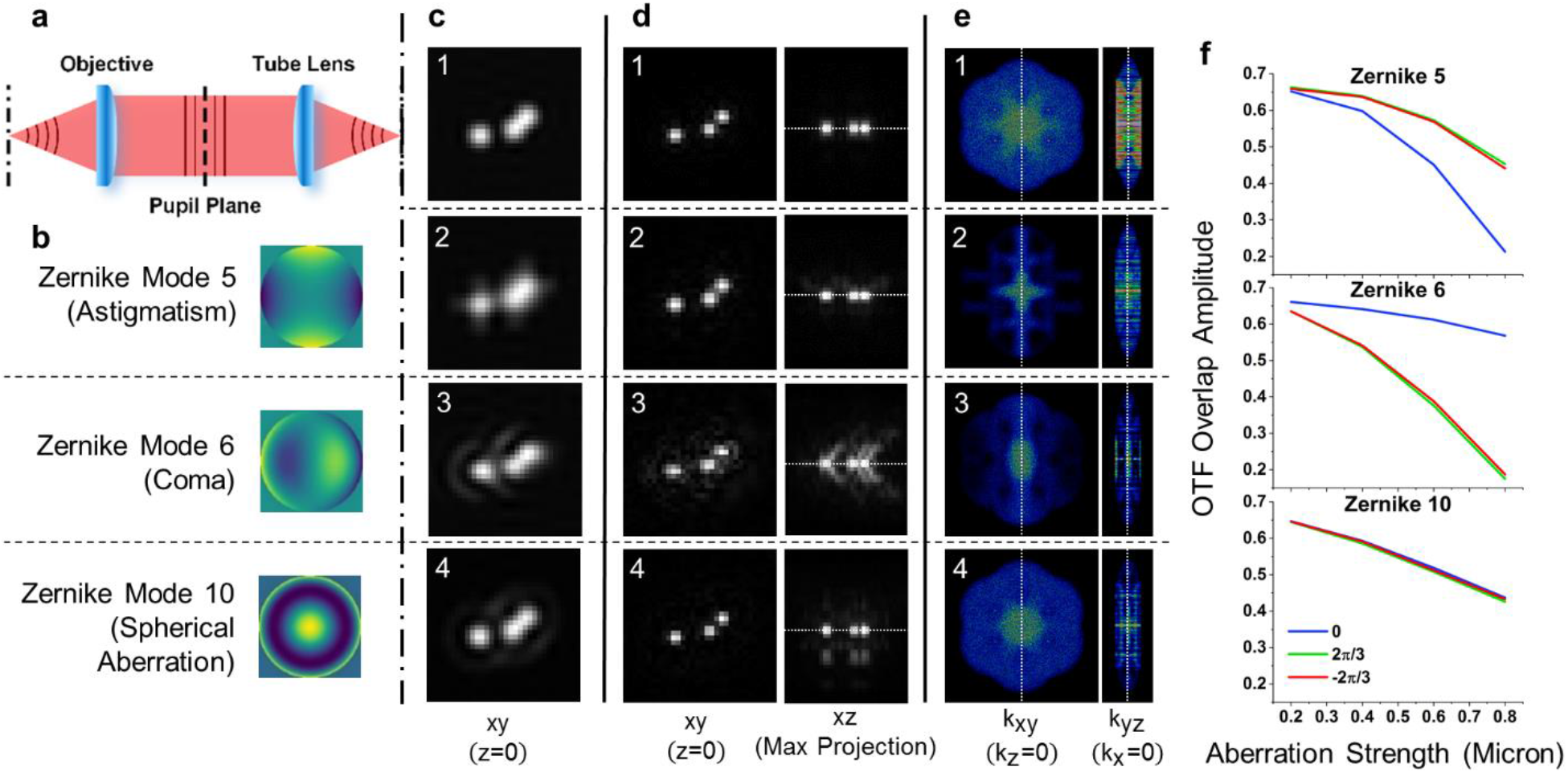
Simulations of the effect of aberrations on 3D-SIM. All spatial images are 2D sections with the third dimension at origin, except for the maximum projection of the 3D-SIM image on XZ plane. The four 3D-SIM effective OTFs are on the same intensity scale. **a,** Schematic illustration of the ideal image formation process with a flat wavefront. **b,** Wavefronts of three typical Zernike modes representing three types of aberrations. **c,** Widefield images of three point objects with different wavefronts at the pupil plane: 1 – flat, 2 – astigmatism, 3 – coma, 4 – spherical aberration. **d,** 3D-SIM image sections. **e,** Effective OTF sections for the 3D-SIM images. **e,** The overlap amplitudes of three pattern orientations (0,2π/3,4π/3) computed by our 3DSIM reconstruction algorithm for the three basic aberration modes (Zernike modes 5, 6 and 10) of increasing strengths. The overlap amplitude is the parameter in the reconstruction algorithm which determines the pattern frequency and strength. The amplitude value reaches its maximum when the position of the shifted data, 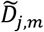. matches the pattern’s wave vector.

Though different aberrations affect the PSF and OTF differently, the strength of the OTF at high frequencies always gets reduced while the low frequencies are much less affected. Therefore we use a spatial frequency based metric function, *M*, defined as [28]:

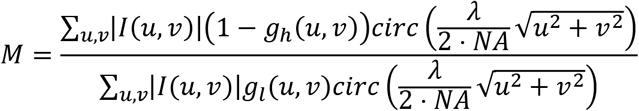

where *g*(*u, v*) is the Gaussian function:

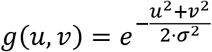

To strengthen metric function’s sensitivity, we choose a slightly larger width for the high frequency filter *g_h_*(*σ* = 64 px) that includes all frequency components higher than *NA/λ* within in the widefield OTF, and a smaller width for the low frequency filter *g_l_*(*σ* = 16 px) that covers frequency components lower than NA/2*λ*. The metric function is calculated on 2D images of size 512 × 512 pixels.

The assessment accuracy of the metric function relies on the image spectrum distribution and the signal to noise ratio (SNR). If the widefield image spectrum is highly concentrated within a limited range, or contains strong high frequency noise, the metric function will lose its efficiency, or even give erroneous results. An ideal object is a sample of sub-resolution fluorescent beads, which are bright and contain all spatial frequencies. However, in biological samples, such objects are rarely present. A customized illumination pattern can be used to optimize the spectrum sampling [43]. As shown in Fig. 5, we illuminated the sample with a dot-array pattern to manually create dot-shaped objects. At the sample each dot has a diameter of 0.75 μm, and the spacing is 0.75 μm.

The AO algorithm operates as follows: Five images are taken for the trial Zernike mode, *Z_l_*, at five different strengths (−2*a*, − *a*, 0, *a*, 2*a*). For each image, the value *M_l_* is calculated from the metric function. A quadratic fit is applied to *x* = (−2*a*, −*a*, 0, *a*, 2*a*) and *y* = *M_l_*. In our experiments, we normally initiated with the step size *a* = 0.1, while for large aberration cases, such as the image of beads under worm body in Fig. 2, we enlarge the step size to *a* = 0.2. From the result, we can locate the peak of the parabola, which is set as the optimal strength for the corresponding Zernike mode. This procedure is repeated on all modes to be corrected to get the optimal strength for each mode separately. Then all modes are combined linearly to form the final corrected wavefront. If needed, the above process can be repeated a set number of times with smaller step size or until the image no longer improves.

**Fig 2.**
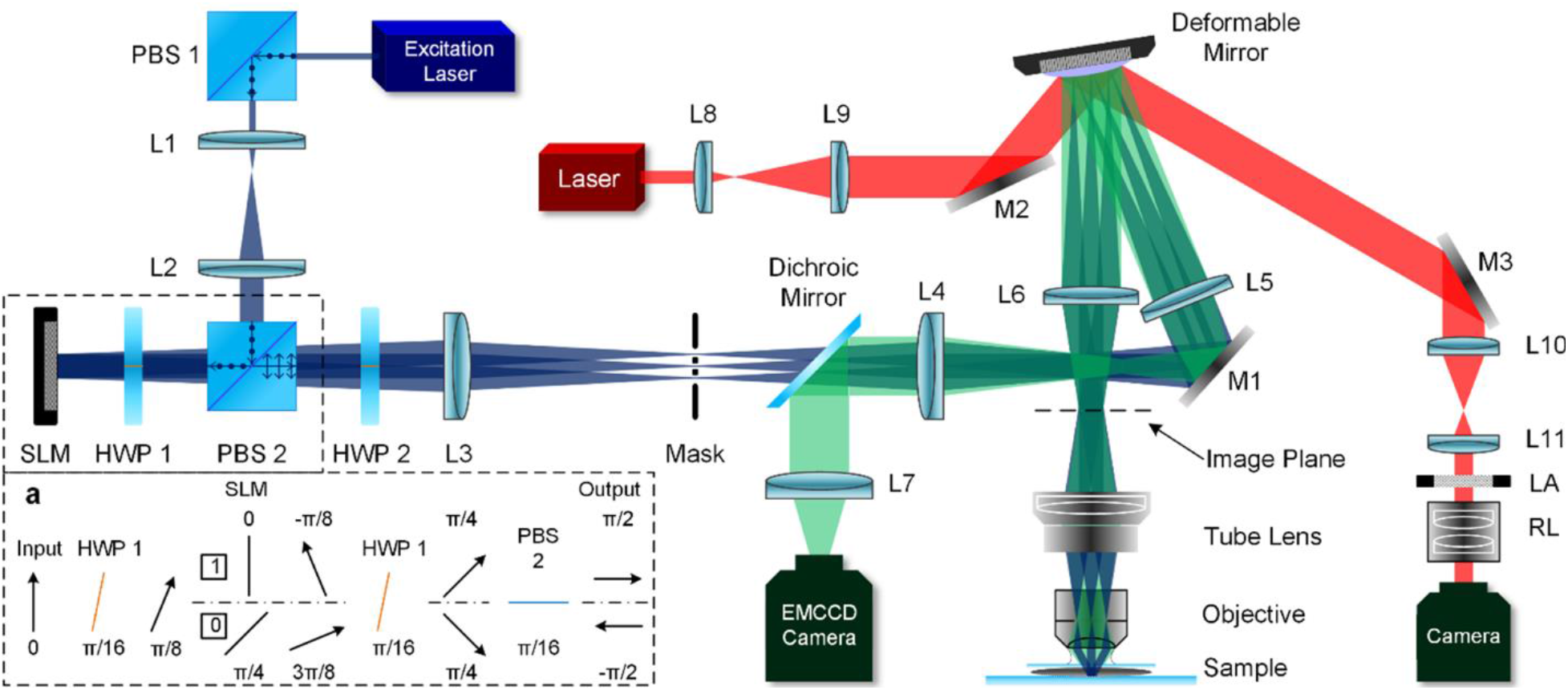
Schematic of the AO-3DSIM microscope setup. **a,** Pattern generation with SLM. SLM: spatial light modulator, HWP: half-wave plate, PBS: polarized beam splitter, LA: lens array, RL: relay lens, M1-M3: flat mirror, L1-L11: lens. f_1_ = 20 mm, f_2_ = 400 mm, f_3_ = 300 mm, f_4_ = 100.7 mm, f_5_ = 250 mm, f_6_ = 250 mm, f_7_ = 300 mm, f_8_ = 30 mm, f_9_ = 300 mm, f_10_ = 350 mm, f_11_ = 170 mm.

Practically, complete correction over a large field of view is rarely achievable, especially on thick biological samples. When the existing aberrations contain anisoplanatic terms, the AO system will fail due to an averaged aberration estimation over the whole widefield image, which could be far from ideal. Therefore, for better performance, a region of interest (ROI) in the image is chosen as the AO target. The correction procedure is then conducted on this target area. This has proven to be effective as shown in Fig. 6, with the disadvantage that the image quality is degraded outside the ROI.

### Experimental Setup

A schematic of the AO-3DSIM system is shown in Fig. 2. We use an Olympus IX71 inverted microscope with a Prior Proscan XY Stage and a Prior 200 micron travel NanoScan Z stage for sample movement and focusing.

Excitation light (Cyan 488 nm - Newport, or 647 nm - Coherent OBIS Lasers) is reflected by a polarizing beam splitter where the s-polarized light (perpendicular to the optical table) is collimated and expanded 20x by a telescope lens set (L1, f_1_ = 20mm, and L2, f_2_ = 400 mm), achieving a quasi-uniform light intensity around the beam center. An iris with a diameter of ~20 mm is inserted after L2 to limit the beam size. The linear polarized beam is sent through a pattern generating unit consisting of a 2048 × 1536 pixel spatial light modulator with pixel size of 8.2 μm (Forth Dimension QXGA-3DM), a polarizing beam splitter cube (10FC16PB.3, Newport) and a half-wave plate. The pattern is generated based on a binary phase grating [13], as shown in Fig. 2a. In this work, we use a biased 15-pixel pattern with 6 pixels ON and 9 pixels OFF. The period at the objective focal plane is 344.08 nm. Thus, in principle, the resolution enhancement with our setup should be 1 + [*λ*/(2 · NA)]/344.08 = 1.81 for red fluorescence (670 nm), and 1.62 for green fluorescence (515 nm). To maximize the modulation depth of the sinusoidal pattern, the polarization state of the illumination light has to be normal to the direction of pattern wave vector for all three angles. This is controlled by a half-wave plate mounted on a fast motorized polarizer rotator (8MRU, Altechna). The dichroic beamsplitter (BrightLine^®^ FF410/504/582/669-Di01, Semrock) used to reflect the illumination beam is designed with low polarization dependence so that the orthogonal relationship between polarization state and pattern wave vector is maintained. A mask on an intermediate pupil plane is used to block all unwanted diffraction orders except for the 0 and ±1 orders. For strengthening the lateral frequency component in the pattern, the diffraction grating is intentionally designed to have higher intensity in the ±1 orders than that in the 0 order. The three beams interfere at the focal plane forming a structured illumination pattern in three dimensions.

The fluorescent emission light from the specimen is collected by the objective (Olympus UPlanSApo 60x water immersion objective with NA of 1.2), and exits the microscope body from the left-side port. A 250-mm lens projects the objective pupil plane onto the DM (ALPAO DM69, 69 actuators and 10mm diameter), which exactly matches the 1.2NA back pupil size of the system. The DM is conjugate to the back pupil plane to provide correction over the whole field of view. The corrected fluorescent image is then directed to the Andor EMCCD camera (DV887DCS-BV with 14bit ADC) through the dichroic beamsplitter. The total magnification of the system is 180 so the effective pixel size of the final image is 89 nm. An emission filter (Semrock BrightLine triple-band bandpass filter, FF01-515/588/700-25) and a notch filter (Semrock StopLine dual-notch filter, NF01-488/647-25×5.0) are put before the camera to block unwanted light, assuring a low background and noise level.

In addition, the system includes a home-built Shack-Hartmann Wavefront Sensor (SHWFS) to measure the DM actuator influence functions and monitor the wavefront applied to the DM. A collimated laser beam with diameter >10 mm is projected onto the DM. The reflected beam is sent through a beam reducing lens pair to image the DM onto the lenslet array (Thorlabs, MLA150-5C). A 1:1 doublet pair (Thorlabs - MAP105050-A) reimages the focal plane of the lenslet array onto the Shack-Hartmann camera (Blackfly BFLY-U3-23S6M).

## Results

We imaged tissue culture cells (α-TN4 lens epithelial cells on a #1.5 glass coverslip) to demonstrate the correction of system aberrations. The imaging was initiated with all the DM actuators set to 0. The size of the 3D image stack is 45.5 *μm* × 45.5 *μm* × 5.2 *μm* in the *x, y*, and *z* dimensions, respectively. The comparison between Fig. 3a and Fig. 3b shows the resolution enhancement from widefield to 3DSIM. And the comparison between Fig. 3b and Fig. 3c shows the effect of AO correction on the 3DSIM image. The Fourier Transform (FT) of the 3DSIM image (Fig. 3d) illustrates the aberrations clearly in the form of a cross shape in each OTF copy, revealing a dominating aberration of astigmatism (Zernike mode 5). Because the system aberrations are isoplanatic, or nearly so, we conducted the AO correction directly on the whole field of view. Fig. 3i shows the wavefront applied on the DM to compensate for the aberrations, and the amplitude of each Zernike mode. As shown in the AO-3DSIM image (Fig. 3c) and its corresponding Fourier transform (Fig. 3e), with AO correction, more fine features show up in the spatial image and the cross shapes disappear from all OTF copies. The improvement in the axial direction can be seen from a comparison among the x-z slices in Fig. 3f (widefield), Fig. 3g (3DSIM), and Fig. 3h (AO-3DSIM). The out-of-focus light, which overwhelms the widefield image, is removed by 3DSIM. And AO correction improves the signal to noise and the fidelity of the image. From the intensity profiles shown in Fig. 3j, we can see the Full Width at Half Maximum (FWHM) getting smaller, the peak value getting larger, and the contrast getting higher from the widefield image without AO to the 3DSIM image with AO in both the lateral and axial directions.

**Fig 3.**
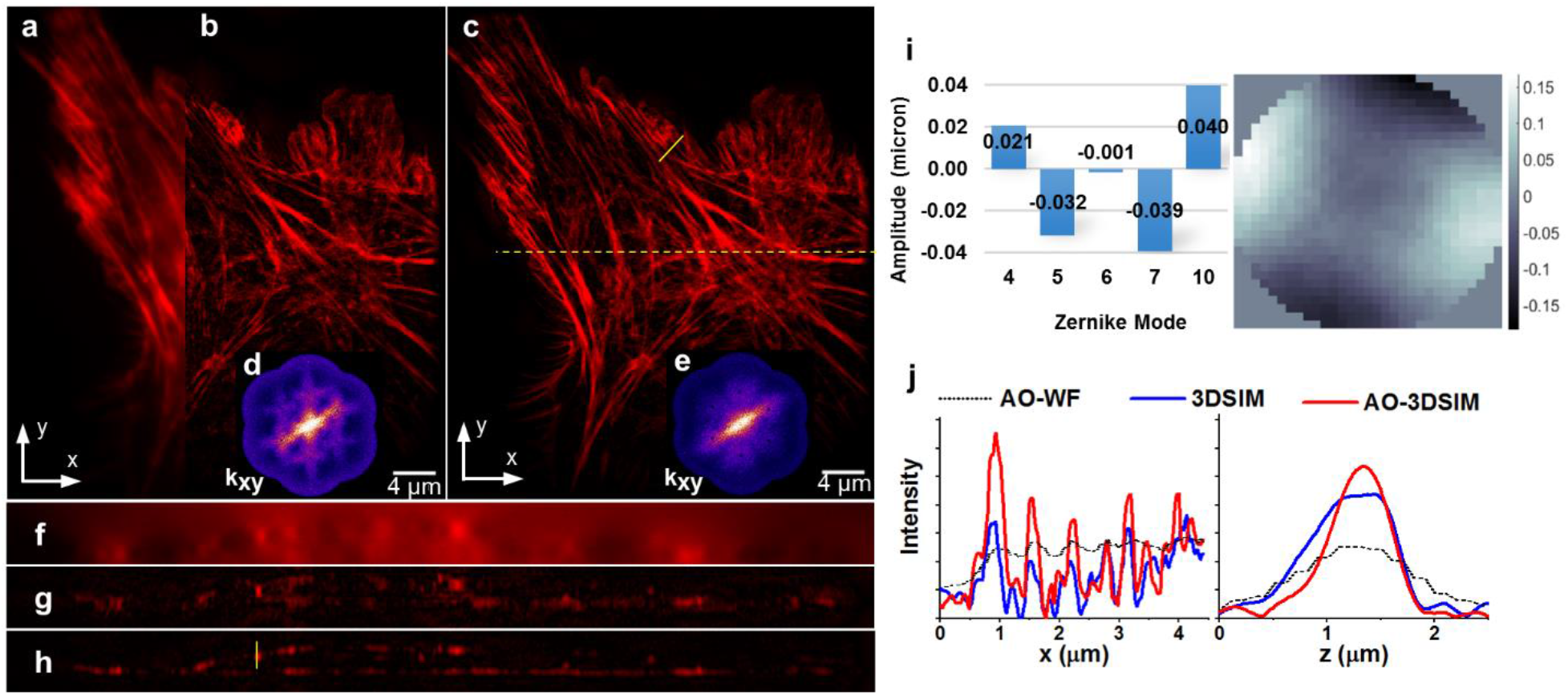
The actin of α-TN4 lens epithelial cell, labelled with Phalloidin-iFluor 647. **a,** Widefield fluorescence image with AO correction. **b,** 3DSIM image without AO correction. **c,** 3DSIM image with AO correction. All three images are x-y slice of the 3D images in the focal plane (z=0). **d,** OTF section at kz=0 of the 3DSIM image without AO correction. **e,** OTF section at kz=0 of the 3DSIM image with AO correction. The x-z slice cut through the dashed yellow line: **f,** widefield with AO; **g,** 3DSIM; **h,** AO-3DSIM. **i,** The amplitudes of Zernike modes and the wavefront posted on the DM. **j.** Intensity profile along the solid yellow lines in x-y and x-z slice images respectively.

We then imaged 100-nm fluorescent beads under a *C. elegans* adult hermaphrodite to demonstrate the correction of sample-induced aberrations. Figs. 4a and 4b illustrate the improvement of 3D resolution and the removal of out-of-focus blur with the application of AO and 3DSIM. Here the imaging is initiated with the DM set to correct system aberrations. Similar to Fig. 3, we can also identify the AO correction from the disappearance of the cross shape in the frequency spectrum shown in Fig. 4a (inset); here, the dominant astigmatism mode is Zernike mode 4, now due to the body of the worm. Figs. 4c and 4d give a more detailed view of the advantages of AO-3DSIM through a close-up look at four 100-nm beads. Without AO correction, we can see artifacts around the beads in all three dimensions. The bead shape is stretched in the lateral plane and elongated along the z-axis. After AO correction, the shape of each bead is recovered and the peak intensity is increased. We see significantly lower noise around the beads, and the intensity distribution along the z-axis is more confined to the focal plane. A quantitative comparison can be seen from the intensity profiles in the x and z directions in Fig. 4e. By Gaussian fitting of the intensity profiles in the x and z directions, and calculating the FWHM, as listed in Table. 1, we get a quantitative evaluation of the improvement due to AO and SIM. We can conclude that the AO-3DSIM system achieves a resolution of 143 nm laterally and 585 nm axially.

**Table 1.** FWHM of 3D image of 100-nm bead

|  | Widefield without AO | Widefield with AO | 3DSIM without AO | 3DSIM with AO |
| --- | --- | --- | --- | --- |
| XY ($\mu\text{m}$) | 0.44144 | 0.416965 | 0.1691 | 0.1428 |
| Z ($\mu\text{m}$) | 1.486 | 1.219 | 1.307 | 0.585 |

**Fig 4.**
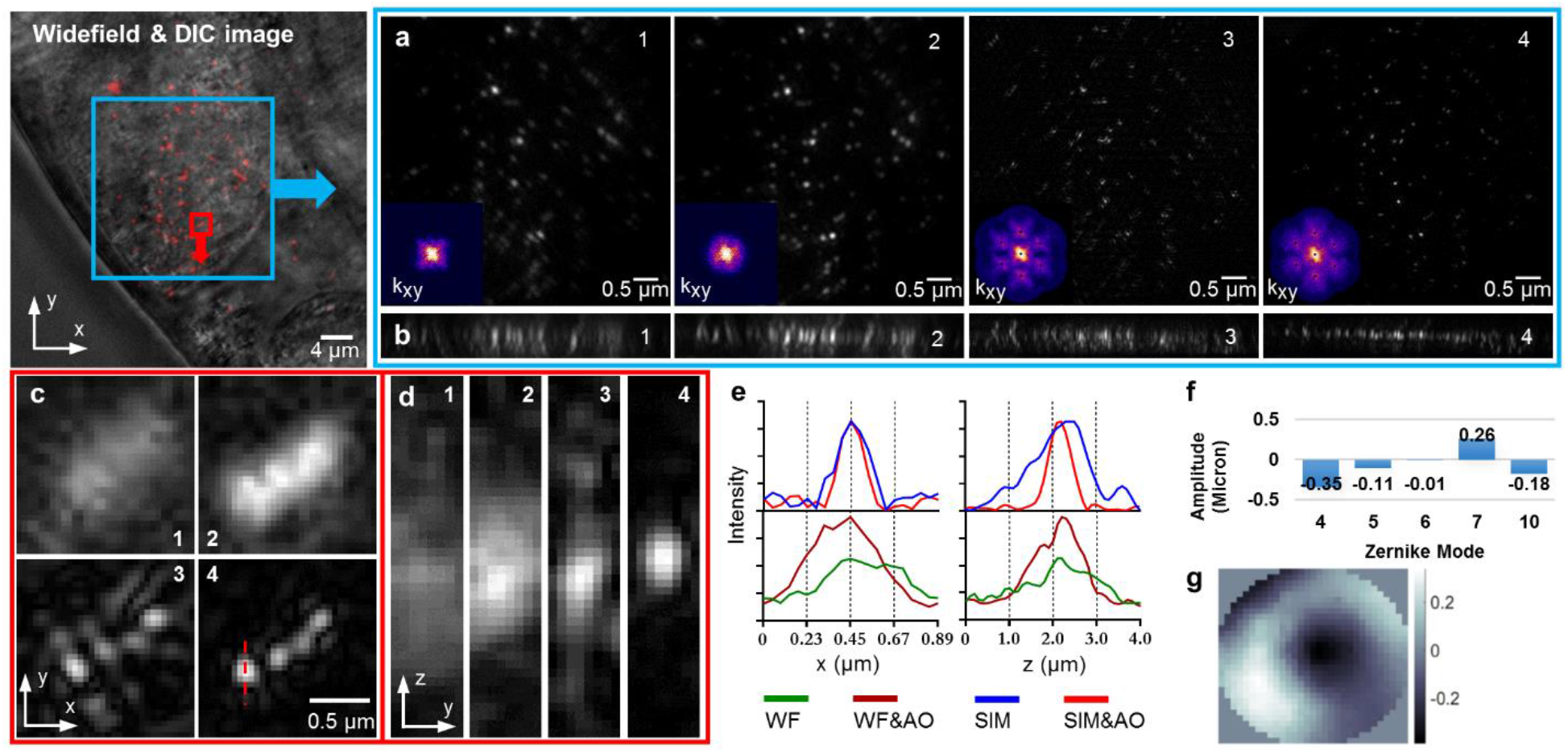
Mixture of 100-nm and 200-nm fluorescent beads under a *C. elegans* worm body. **a,** In-focus x-y section images of area within blue square, and their corresponding FT. **b,** Maximum intensity projection of x-z sections. **c,** Zoom in of the area within the red square. **d,** The x-z slice cut through the dashed red line. **e,** Intensity profile. **f,** The amplitudes of Zernike modes applied on the DM. **g.** The wavefront applied to the DM. (**1,** Widefield image without AO correction. **2,** Widefield image with AO correction. **3,** 3DSIM image without AO correction. **4,** 3DSIM image with AO correction.)

Next, we applied AO-3DSIM on a more challenging object: GFP fused to the protein TBH-1 expressed in the RIC neurons in *C. elegans* [48]. Tyramine beta-hydroxylase (TBH-1) is an enzyme involved in the octapamine biosynthetic process. Here, we focus on the axon shown in Fig. 5. The challenge arises from the assessment of the image quality using our metric function. The frequency spectrum of the axon image is heavily biased along one direction which prevents the frequency-based metric function from working optimally. And the GFP fluorescence cannot survive too many AO correction loops. So, for this sample, we used dot pattern illumination to boost the frequency distribution in Fourier space while conducting AO corrections [43]. The dot illumination pattern is shown in Fig. 5a; it creates more high frequencies in Fourier space as indicated by the arrows in Fig. 5a (image spectrum). The AO correction is done successfully within one AO correction loop (25 images in total: 5 Zernike modes, 5 trial amplitudes for each mode). Fig. 5e shows the image assessment results for Zernike modes 4 to 7, and 10. The effect of AO correction can be seen by comparing Figs. 5a and 5b in both the spatial domain and the frequency domain. The filament structure is clearer and the signal intensity becomes stronger after AO correction. The image spectrum contains more high frequency components after correction. We took a 4 *μm* stack in z axis with a step of 0.1 μm for 3DSIM. With AO correction, the image of the nerve fiber, Fig. 5d, has better resolution and more fidelity compared to the dim and noisy image without AO correction, Fig. 5c. In x-z slices, the peak intensity after AO correction is higher than before AO correction, while the noise level is much lower. The isolated nerve fibers can be clearly recognized in the image.

**Fig 5.**
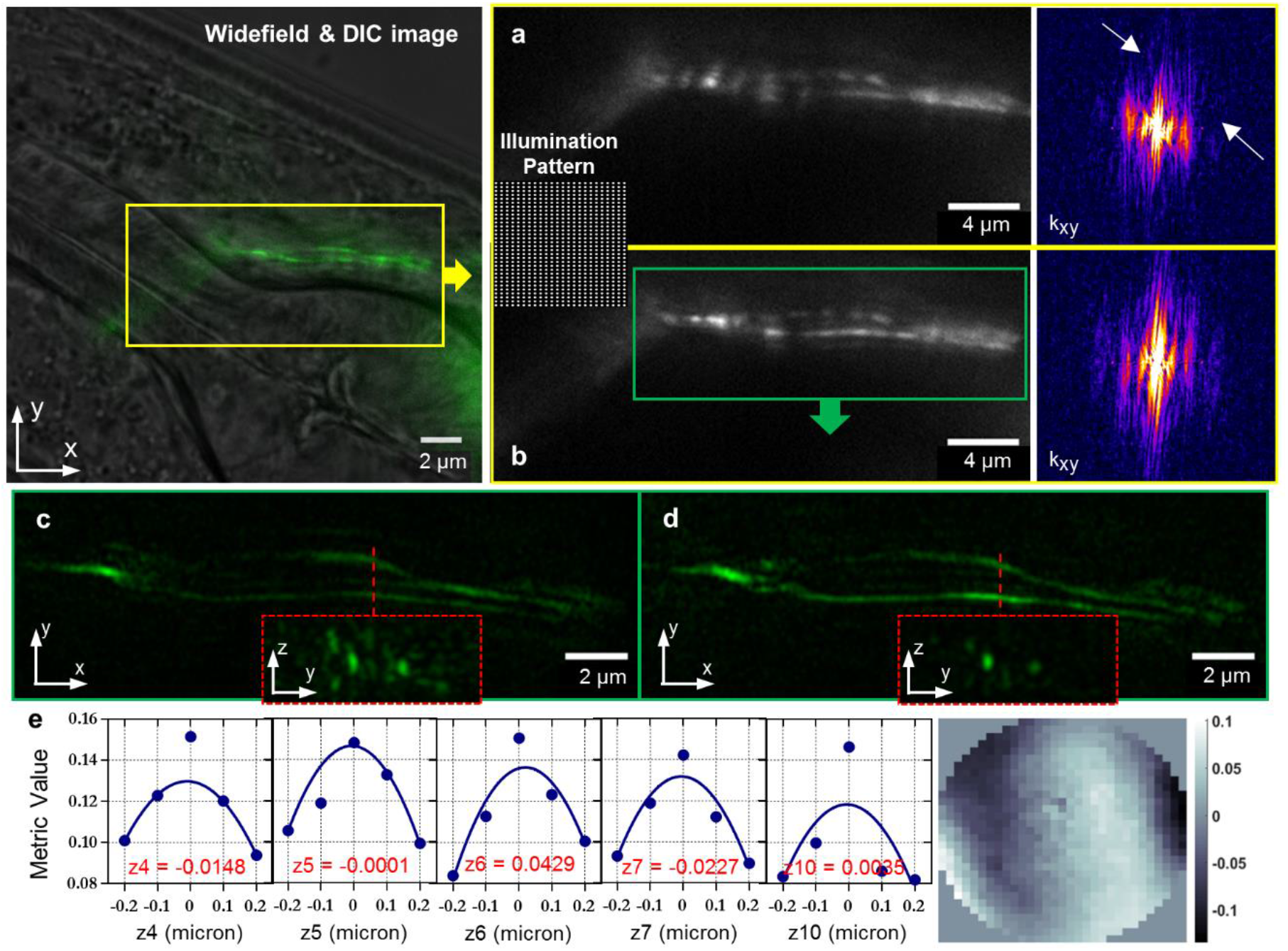
Live *C. elegans* with TBH-1::GFP expressed in RIC interneurons in the lateral ganglion. **a,** Dot pattern illuminated axon and its Fourier transform before AO correction. **b,** Dot pattern illuminated axon and its Fourier transform after AO correction. **c,** 3DSIM image of axon before AO correction and the y-z slice cut through the dashed red line. **d,** 3DSIM image of axon after AO correction and the y-z slice cut through the dashed red line. **e,** The metric function values and polynomial fitting curves of each trial aberration modes. The amplitude of each mode is the peak location of each curve. **f,** The wavefront posted on the DM.

Last, but not least, we imaged the endoplasmic reticulum (ER) of the filamentous fungus *Magnaporthe oryzae*, also known as rice blast fungus, growing inside rice plant cells. The fungus was transformed to constitutively express GFP fused with an N-terminal secretory signal peptide and a C-terminal ER retention signal peptide, which labels the ER network [49]. For live-cell imaging, we used hand-cut optically clear leaf sheath tissue (~60 μm thick), consisting of a layer of epidermal cells, which the fungus colonized, and a few underlying layers of mesophyll cells. As shown in Fig 6a, fungal hyphal cells reside within a rice epidermal cell. The green dash-dotted line in Fig. 6a marks a rice cell wall of the mesophyll cells. Due to the spatial variance over the whole field of view, we cropped the image and conducted AO correction upon the area inside the white dotted square. With AO correction and SIM, we can clearly identify the integral structure of ER and distribution around the nuclei in all three dimensions as shown in Fig. 6d. While, in Fig. 6c, due to the aberration and its induced reconstruction artifacts, the image looks fuzzy and the ER structure is not smooth and continuous as it is in Fig. 6d. The difference is also seen in Fourier space. Comparing the FT of the image without correction in Fig. 6e, the image FT after correction in Fig. 6f has stronger mid and high frequency components, revealing that AO correction restores information and recovers image quality.

**Fig 6.**
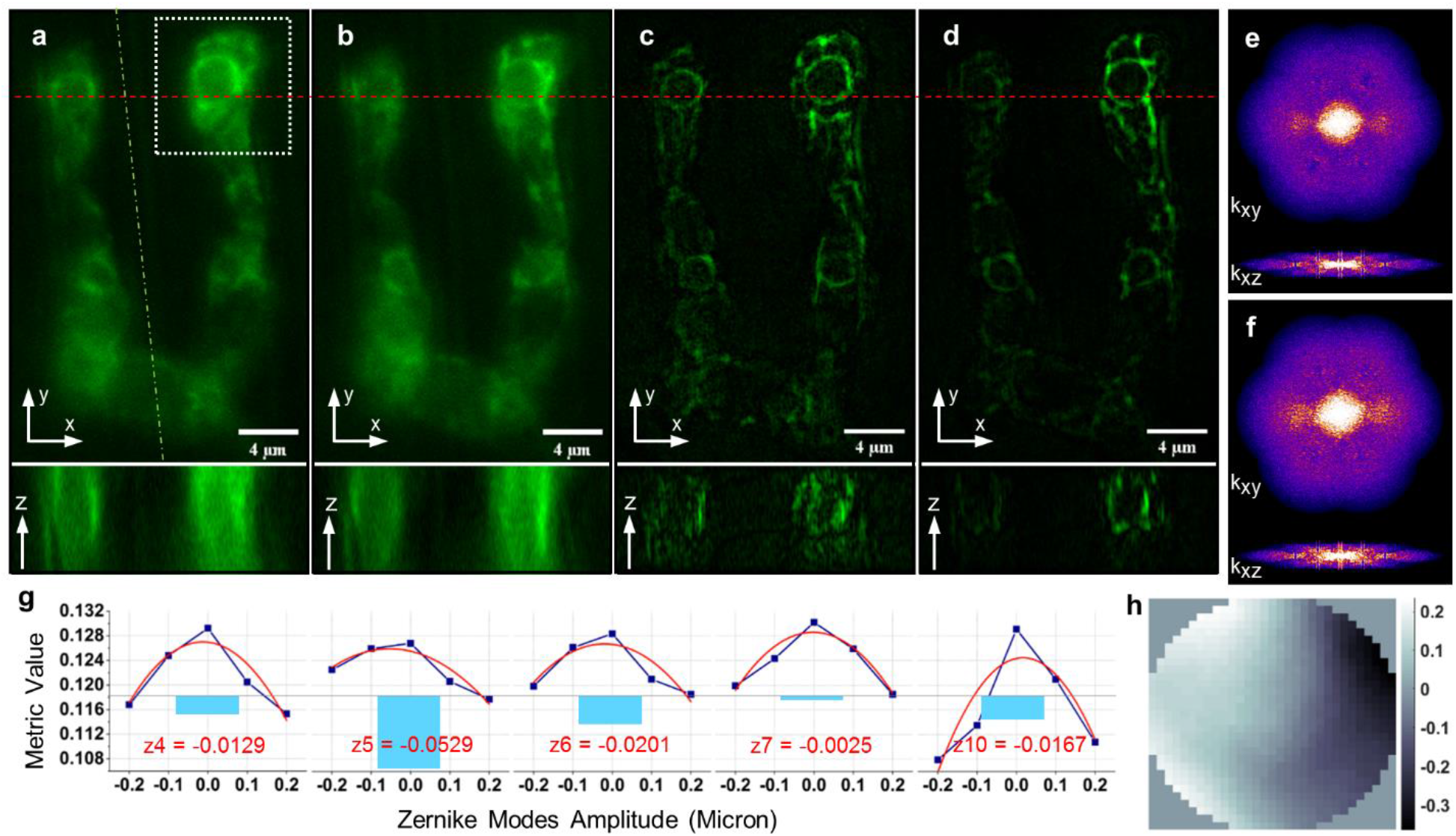
GFP-labelled endoplasmic reticulum of live *M. oryzae* hyphal cells growing inside rice plant cells. XY image section in focus and XZ image section along the red dash line: **a,** Widefield image with only system aberration corrected. **b,** Widefield image with both sample and system aberrations corrected. **c,** 3DSIM image with only system aberration corrected. **d,** 3DSIM image with both sample and system aberrations corrected. **e,** Effective OTF of 3D-SIM image without sample aberration correction. **f,** Effective OTF of 3D-SIM image with sample aberration correction. **g,** The metric function values of the sample aberration correction process and polynomial fitting curves of each trial aberration modes. The red number is the amplitude of each mode. **h,** The wavefront applied to the DM.

To evaluate the overall performance of AO-3DSIM, image SNR is calculated. The noise is defined as the standard deviation (STD) around the maximum, which is defined as the signal. The image SNR results are in Table. 2 showing the improvement due to the AO correction.

**Table 2.** SNR Results

| Sample | Exposure Time (s) | EMCCD Gain | SNR |  |  |  |
| --- | --- | --- | --- | --- | --- | --- |
|  |  |  | Widefield |  | 3D-SIM |  |
|  |  |  | Before AO | After AO | Before AO | After AO |
| Actin | 0.1 | 150 | 35.04 | 50.11 | 20.63 | 33.48 |
| Beads | 0.1 | 200 | 34.08 | 53.66 | 52.49 | 177.91 |
| Axon | 0.05 | 100 | 10.33 | 10.51 | 28.71 | 39.74 |
| ER | 0.1 | 150 | 36.79 | 45.39 | 14.23 | 20.33 |

## Discussion and Conclusion

In conclusion, we have presented a three-dimensional super-resolution fluorescence microscopy system based on a combination of three-dimensional structured illumination and adaptive optics. We have demonstrated a lateral resolution of 140nm and an axial resolution of 585 nm at 670 nm emission wavelength when imaging through the body of a *C. elegans* roundworm, and we have shown that the image-based AO correction is capable of correcting large aberrations and restoring images that were badly distorted by optical aberrations. Under severe conditions, such as imaging neuron filaments in *C. elegans*, patterned illumination can be used to support the image quality assessment. Moreover, the AO correction is also an enhancement of the illumination pattern projected on the sample, and results in an improved final SR image reconstruction.

AO-3DSIM is a promising method to acquire three-dimensional super-resolution images in live thick samples with improved fidelity. Several areas of biology will benefit from the improved resolution in living samples including imaging of *C. elegans*, plant roots, and tissue spheroids which are important for three-dimensional cell culture. AO-3DSIM has enormous potential to help understand sub-cellular dynamics in organisms and tissues *in vivo.*

## Acknowledgments

We thank James D. Lauderdale for help with tissue culture.

## Author Contributions

P.K devised and supervised the project. R.L. and P.K. built the microscope and wrote the software. R.L. prepared the tissue culture samples, acquired and analyzed all the data. E.T.K. provided expertise and assistance with *C. elegans* biology and imaging, and provided some of the *C. elegans* strains. J.Z. and C.H.K. prepared the *M. oryzae* samples and provided expertise and assistance with *M. oryzae* biology and imaging. R.L. and P.K. wrote the manuscript with assistance from all authors.

## Materials and Correspondence

Please address all correspondence and material requests to Peter Kner at.

## Sample Preparation

1. Beads under *C. elegans* worms. The diluted 100-nm fluorescent beads (T7279, Life Technologies, TetraSpeck^™^ Microspheres, 0.1 μm, fluorescent blue/green/orange/dark red) were dried onto charged slides (9951APLUS-006, Aqua ColorFrost Plus). Wild-type C. elegans worms were placed above the beads with 10 μL of a 50 mM solution of phosphatase inhibitor Tetramisole hydrochloride (T1512, Sigma-Aldrich) to inhibit worm movement. Before the Tetramisole solution dries completely, 10 μL of glycerol was put on the slide, and the coverslip was mounted and fixed carefully.
2. Cell culture. α-TN4 lens epithelial cells were grown on poly-L-lysine coated coverslips, fixed with 4% paraformaldehyde in Phosphate-buffered saline (PBS) pH ~7.4. And then the fixed cells were incubated in 1 mL Phalloidin-iFluor 647 (ab176759) solution (1 μL 1000x stock solution in 1 mL PBS) under 4°C overnight to stain the F-actin. After staining, the cells were washed with PBS three times, and mounted with antifade medium (VECTASHIELD^®^ H-1000) for imaging.
3. *C. elegans.* The *C. elegans* strain MT9971 (*nIs107* [*tbh-1*::GFP + *lin-15(+)*] III.) was obtained from the Caenorhabditis Genetics Center. The worms were grown on 3x nematode growth media (NGM) agar plates (3 g/L NaCl, 7.5 g/L bacto peptone, 20 g/L agar, 1 mM MgSO4, 1 mM CaCl2, 5 mg cholesterol in ethanol, and 25 mM K2HPO4) and fed OP50 bacteria that was grown on the agar plates. To mount worms: a drop of 2% agarose was placed onto a clean slide, and then covered with another clean slide to flatten the agarose. After the agarose becomes solid, the top slide is then gently shifted to separate the two slides, allowing the agarose pad to adhere to one of the two slides. 10 μL of 20 mM sodium Azide (NaN3) and 10 μL of 50 mM Tetramisole solution, both dissolved in M9 buffer (3.0 g/L KH2PO4, 6.0 g/L Na2HPO4, 0.5 g/L NaCl, 1.0 g/L NH4Cl), were placed on the center of the agarose pad. The worm to be observed was then transfer into the drop with a worm pick. A clean coverslip was gently put on and sealed with nail polish.
4. *M. oryzae* growing inside rice cells. *M. oryzae* transgenic strain CKF4019, expressing GFP retained in the ER lumen, was generated by transforming *M. oryzae* wild-type strain O-137 with plasmid pCK1724 [49]. The plasmid carries GFP fused at 5’end with the sequence encoding the BAS4 signal peptide and at 3’ end with the sequence encoding the ER retention signal peptide HDEL. A rice sheath inoculation method was used to prepare optically clear rice tissue with the fungus as previously described [49]. Briefly, rice sheaths from 20-day old rice cultivar YT16 were excised to about 8 cm in length, and subsequently the inner epidermal tissue of the excised sheath was inoculated with CKF4019 conidial suspension (1×10^5^ conidia/ml in water). At 30 hours post inoculation, the inoculated rice sheath was hand-trimmed with a razor blade to produce optically clear leaf sheath slices with 3-4 cell layers deep and about 60 μm thick. The trimmed sheath was mounted on a slide in water under a coverslip for live-cell imaging.

